# Host responses in an *ex-vivo* human skin model challenged with *Malassezia sympodialis*

**DOI:** 10.1101/2020.07.22.215368

**Authors:** Dora E. Corzo Leon, Donna M. MacCallum, Carol A. Munro

**Affiliations:** School of Medicine, Medical Sciences and Nutrition, Institute of Medical Sciences, University of Aberdeen, AB25 2ZD, United Kingdom

**Keywords:** *Malassezia sympodialis*, skin model, immune response, cytokines, AMPs

## Abstract

*Malassezia* species are a major part of the normal mycobiota and colonise mainly sebum-rich skin regions of the body. This group of fungi cause a variety of infections such as pityriasis versicolour, folliculitis and fungaemia. In particular, *Malassezia sympodialis* and its allergens have been associated with non-infective inflammatory diseases such as seborrheic dermatitis and atopic eczema. The aim of this study was to investigate the host response to *M. sympodialis* on oily skin (supplemented with oleic acid) and non-oily skin using an *ex-vivo* human skin model. Host-pathogen interactions were analysed by SEM, histology, gene expression, immunoassays and dual species proteomics. The skin response to *M. sympodialis* was characterised by increased expression of the genes encoding β-defensin 3 and RNase7, and by high levels of S100 proteins in tissue. Supplementation of oleic acid onto skin was associated with direct contact of yeasts with keratinocytes and epidermal damage. In oily conditions, skin response to *M. sympodialis* showed no gene expression of AMPs, but increased expression of *IL18*. In supernatants from inoculated skin plus oleic acid, TNFα levels were decreased and IL-18 levels were significantly increased.

## INTRODUCTION

The genus *Malassezia*, previously known as *Pityrosporum*, is a group of lipophilic yeasts. *Malassezia* species are part of the normal mycobiota and colonise several regions of the body, mainly sebum-rich skin areas such as the scalp and thorax (Marcon & Powell, 1992). To date, 17 *Malassezia* species have been proposed (Sparber & LeibundGut-Landmann, 2017). *Malassezia* spp. are the most abundant genus on the skin of individuals with psoriasis and atopic eczema (Zhang *et al*, 2011; Paulino *et al*, 2008; Paulino LC *et al*, 2006) and are highly increased in seborrheic dermatitis and dandruff (Park *et al*, 2012; DeAngelis *et al* 2005).

*M. globosa, M. furfur, M. restricta* and *M. sympodialis* are the most frequent *Malassezia* species on both healthy (Findley *et al*, 2013) and diseased skin, such as pityriasis versicolor and seborrheic dermatitis (Prohic & Ozegovic, 2007; Prohic, 2010; Amado *et al*, 2013; Lyakhovitsky *et al*, 2013; Lian *et al*, 2014; Rodoplu *et al*, 2014). *M. sympodialis* is the most frequent coloniser of healthy skin (Findley *et al*, 2013), but is found less frequently in *Malassezia* related-diseases, such as seborrheic dermatitis and pityriasis versicolour (Prohic & Ozegovic, 2007; Prohic, 2010; Amado *et al*, 2013).

Most *Malassezia* species are unable to synthesise fatty acids and degrade carbohydrates and are dependent upon the acquisition of exogenous fatty acids. *Malassezia* species have a large repertoire of lipolytic enzymes such as lipases, phospholipases and esterases (Sparber & LeibundGut-Landmann, 2017; Saunders *et al*, 2012). Lipases hydrolyse sebum triglycerides from the host skin to release fatty acids (oleic acid and arachidonic acid). Lipase activity is significantly higher in *M. globosa* and *M. pachydermatis* than in *M. sympodialis* and *M. slooffiae*, although normal phospholipase activity is found in these species (Juntachai *et al*, 2009). The fatty acids oleic acid (OA) and arachidonic acid are released by phospholipase and lipase activity and have an irritating/inflammatory effect on skin (Ashbee & Evans, 2002). OA is also increased in dandruff scalps compared with non-dandruff scalps (Jourdain *et al*, 2016) and increases the severity of flaking (or *stratum corneum* desquamation) facilitating better penetration of *M. sympodialis* so that the fungus directly interacts with cells of the inner skin layers (DeAngelis *et al*, 2005).

Exogenously acquired fatty acids contribute to the formation of a thick cell wall in *Malassezia* sp. characterised by a unique lipid-rich outer layer, which contributes to triggering the immune response against this group of fungi (Thomas *et al*, 2008; Gioti *et al*, 2013b). Human studies have shown that cytokine levels (IL-1α, IL-1β, IL-2, IL-4, IFN-γ, IL-10 and IL-12) in the skin of individuals with seborrheic dermatitis and *Malassezia* folliculitis were higher than levels in the skin of healthy volunteers (Faergemann *et al*, 2001). However, this pattern of cytokine induction varied depending on the fungal cell wall structure. Higher levels of IL-8 and lower levels of IL-10 were produced by keratinocytes *in vitro* when they were stimulated with *M. sympodialis* lacking the lipid-rich outer layer compared to the same yeasts with the outer layer (Thomas *et al*, 2008).

Antimicrobial peptides (AMPs) are key in the innate immune response to *Malassezia* spp. Individuals with pityriasis versicolour have significantly higher AMP levels (β-defensin 2, β-defensin 3, S100A7 and RNase7) in their skin (Brasch *et al*, 2014). The β-defensins are specifically increased in the *stratum corneum*, RNase7 in the *stratum granulosum*, and S100A7 in the *stratum corneum, granulosum* and *spinosum* (Brasch *et al*, 2014). *Malassezia* sp. also play a role in NLRP3 inflammasome activation when yeast cells are sensed by dendritic cells but not by keratinocytes (Kistowska *et al*, 2014). Activation of NLRP3 depends on Dectin-1 and leads to high expression of caspase-1-dependent IL-1β in dendritic cells of patients with seborrheic dermatitis (Kistowska *et al*, 2014). The adaptive immune response against *Malassezia* is characterised by the production of specific IgG and IgM antibodies in healthy individuals and specific IgE antibodies in atopic eczema (Glatz *et al*, 2015).

Atopic eczema (AE) is a chronic inflammatory disease affecting up to 20% of children and 3% of adults (Nutten, 2015). Multiple factors have been associated with AE, such as impairment of skin barrier function due to physical (scratching, skin dryness) or chemical (pH changes due to soaps) damage, genetic factors (mutations in *FLG* and *SPINK5* genes encoding filaggrin and serine protease inhibitor Kazal-type 5 protein, respectively), and environmental factors (cold climate, no breastfeeding, pollution) (Nutten, 2015; Sääf *et al*, 2008). *Malassezia* sp. has been linked with AE pathogenesis as *Malassezia* sp. allergens induce specific IgE antibodies and autoreactive T-cells that can cross-react with skin cells (Glatz *et al*, 2015; Zargari *et al*, 2001).

The *M. sympodialis* genome contains 13 allergen genes, Mala S1, Mala S5 to S13 and three orthologs of *M. furfur* allergens (Mala F2, 3, and 4) (Gioti *et al*, 2013; Andersson *et al*, 2003). Some of these allergens are highly similar to human proteins (Mala S11, Mala S13) and have been specifically linked to cross-reactive immune responses (Gioti *et al*, 2013; Schmid-Grendelmeier *et al*, 2005; Roesner *et al*, 2019).

The aim of this study was to investigate the host skin response to *Malassezia sympodialis.* An *ex-vivo* human skin model was used either directly for non-oily skin or supplemented with oleic acid to represent oily skin. Host-pathogen interactions were analysed by SEM, histology, gene expression, immunoassays and proteomics.

The skin response to *M. sympodialis* was characterised by increased expression of the genes encoding β-defensin 3, and RNase7 and high levels of S100 proteins. Supplementation of the skin with oleic acid resulted in epidermal damage, direct contact of yeasts with keratinocytes, and AMP gene expression was not detected. TNFα levels were decreased in supernatants from inoculated skin plus oleic acid. IL-18 levels were significantly increased in the same supernatants and *IL18* gene expression was increased in tissue.

## MATERIALS AND METHODS

### An *ex-vivo* human skin model

The *ex-vivo* human skin model was set up as previously described with modifications (Corzo-León *et al*, 2019). Human skin tissue (no stretch marks), without an adipose layer (adipose layer removed by the surgeon during surgery), from abdominal or breast surgeries was supplied by Tissue Solutions^®^ Ltd. (Glasgow, UK). Human tissue from four different donors was obtained according to the legal and ethical requirements of the country of collection, with ethical approval and anonymous consent of the donor or nearest relative. Tissue Solutions^®^ comply with the UK Human Tissue Authority (HTA) on the importation of tissues. Explants were transported at 4 °C with cooling packs and maintained at 4 °C until processed, which occurred within 36 h of surgery.

Skin was washed with DMEM (supplemented with 1% v/v antibiotics (penicillin and streptomycin) and 10% heat inactivated FBS, Thermo-Fisher Scientific, Loughborough, UK) and kept moist in a Petri dish in the same medium. The explant was cut into 1 cm^2^ pieces. The surface of each piece was gently wounded using a needle, without penetrating the entire skin thickness, by pricking with a needle several times (6-10) to a depth of approximately 2-3 mm.

After wounding, each piece of skin was placed into an individual well of a 6-well plate. An air-liquid interphase was maintained by adding 1 ml of supplemented DMEM. A well containing only DMEM growth medium was added as a negative control and served as a contamination control. The medium was changed every 24 h and the spent medium was stored in 2 ml tubes for subsequent analysis at the same time points as the rest of the samples. Recovered medium was stored at -80 °C until analysed. In a previous study (Corzo-León *et al*, 2019) the skins samples were demonstrated to remain viable for 14 days under these conditions using the TUNEL system.

Skin explants were inoculated by applying 10 µl of fungal suspension (1×10^6^ yeasts) directly on to the epidermis. Yeast suspension was prepared as described below and resuspended in PBS. Uninfected/non-inoculated skin controls were included in all experiments. Two additional experimental conditions were added: 1) uninfected skin with 10 µl 100% oleic acid (Thermo-Fisher Scientific) applied to the surface of the skin explant and 2) infected skin with same *M. sympodialis* inoculum, followed by application of 10 µl 100% oleic acid on to the surface of the skin explant.

Skin samples were incubated for six days at 37 °C and 5% CO_2_ before being recovered in a Petri dish. Prior to processing, the macroscopic appearance of explants was evaluated by eye and images captured with a Stemi 2000-c Stereo Microscope (Carl Zeiss, Oberkochen, Germany). Samples were then processed depending for further analyses.

Tissue samples for histology were placed into moulds, embedded in OCT compound (Cellpath Ltd. Newtown, UK) and flash-frozen with dry ice and isopentane. These samples were stored at -20 °C for immediate analysis or at -80 °C for longer term storage. For scanning electron microscopy (SEM), tissue samples were fixed in glutaraldehyde buffer (2.5% glutaraldehyde in 0.1 M cacodylate) overnight at 4 °C and sent to the Microscopy and Histology technology hub, University of Aberdeen, for further sample preparation. Tissue samples for RNA extraction were cut into smaller pieces and placed in a microcentrifuge tube with RNAlater^®^ (Sigma, Dorset UK) for subsequent RNA extraction. These samples were stored at -20 °C for immediate analysis or at -80 °C for longer term storage. Experiments were replicated at least three times using skin from different human donors.

### Fungal strains and culture conditions

Modified Dixon (mDixon) broth and agar (3.6% w/v malt extract, 2% w/v desiccated ox-bile, 0.6% w/v bacto tryptone, 0.2% v/v oleic acid, 1% v/v Tween 40, 0.2% v/v glycerol, 2% w/v bacto agar for agar plates) was used to grow *Malassezia sympodialis* (ATCC 42132) for skin experiments. Yeast cells were grown on mDixon agar at 35 °C for 4-5 days in a static incubator, one colony was selected from an agar plate and inoculated into 10 ml of mDixon broth for 4 days at 37 °C in a shaking incubator at 200 rpm. Yeast cells were recovered and a final inoculum of 1×10^6^ yeasts in 10 µl of PBS (1×10^8^/ml) was prepared and applied to the skin surface.

### Scanning Electron Microscopy (SEM) and Histopathology

Several microscopic analyses were performed on recovered skin tissue for histological confirmation of fungal infection. Sections (0.6 µm) were cut from the frozen OCT blocks for histological analysis and stained with fluorescent dyes (1 µg/ml calcofluor white (CFW), and propidium iodide (1 µg/ml). Images were captured with a Deltavision™ confocal microscope (GE Healthcare, Buckinghamshire UK). SEM samples were observed using a Zeiss™ EVO MA10 Scanning Electron Microscope. Images were generated by detection of secondary electrons (SE1) and backscatter electron detection (NTS BSD) and captured at 10 kV resolution and at different magnifications.

### Gene expression and proteomics analyses

Tissue samples for RNA extraction were thawed and the RNAlater^®^ discarded before processing the samples. Samples were washed twice with PBS. RNA and proteins were extracted from the same recovered samples in a single two-day sequential process based on previously published methods (Chomczynski & Sacchi, 1987; Berglund *et al*, 2007; Corzo-León *et al*, 2019).

The RNA yield and purity were evaluated by Nanodrop™ spectrophotometry (Thermo-Fisher Scientific) and samples were stored at -80 °C until further use. All samples had an initial yield between 200 and 800 ng/µl and a 260/280 ratio between 1.8 and 2.0. To produce cDNA, RNA samples (1 µg) were treated with DNase I (1 U/µl per 1 µg of RNA sample) (Thermo-Fisher Scientific), then reverse transcription carried out using the SuperScript™ IV first-strand synthesis system (Thermo-Fisher Scientific) with Oligo dT primer, following the manufacturer’s instructions.

Intron spanning primers and qRT-PCR assays were designed using Roche’s Universal Probe Library Assay Design Centre (lifescience.roche.com/en_gb/brands/universal-probe-library.html#assay-design-center) for different target genes known to be expressed in skin. Genes, accession numbers, Roche probes paired with target primers and primer sequences are shown in Table 1. The Roche probes were hydrolysis probes labelled at the 5□ end with fluorescein (FAM) and at the 3□ end with a dark quencher dye. A reference gene (*B2M* encoding β2-microglobulin) primer pair and probe was also designed (Lossos *et al*, 2003) using the Eurogentec web tool (secure.eurogentec.com/life-science.html) (Table 1). The probe was modified at the 5□ end with Cy5 and at the 3□ end with quencher QXL 670. qRT-PCR reactions (10 µl) were set up in Light cycler 480 plates using the LightCycler 480 probe master mix (Roche, Welwyn Garden City UK) according to the manufacturer’s instructions (Tables 2). Dual hydrolysis probe assays were analysed in the same well, FAM probe was used for target gene primer pairs and Cy5 probe for the reference gene. For each cDNA, assays were performed in triplicate.

**Table 1.**
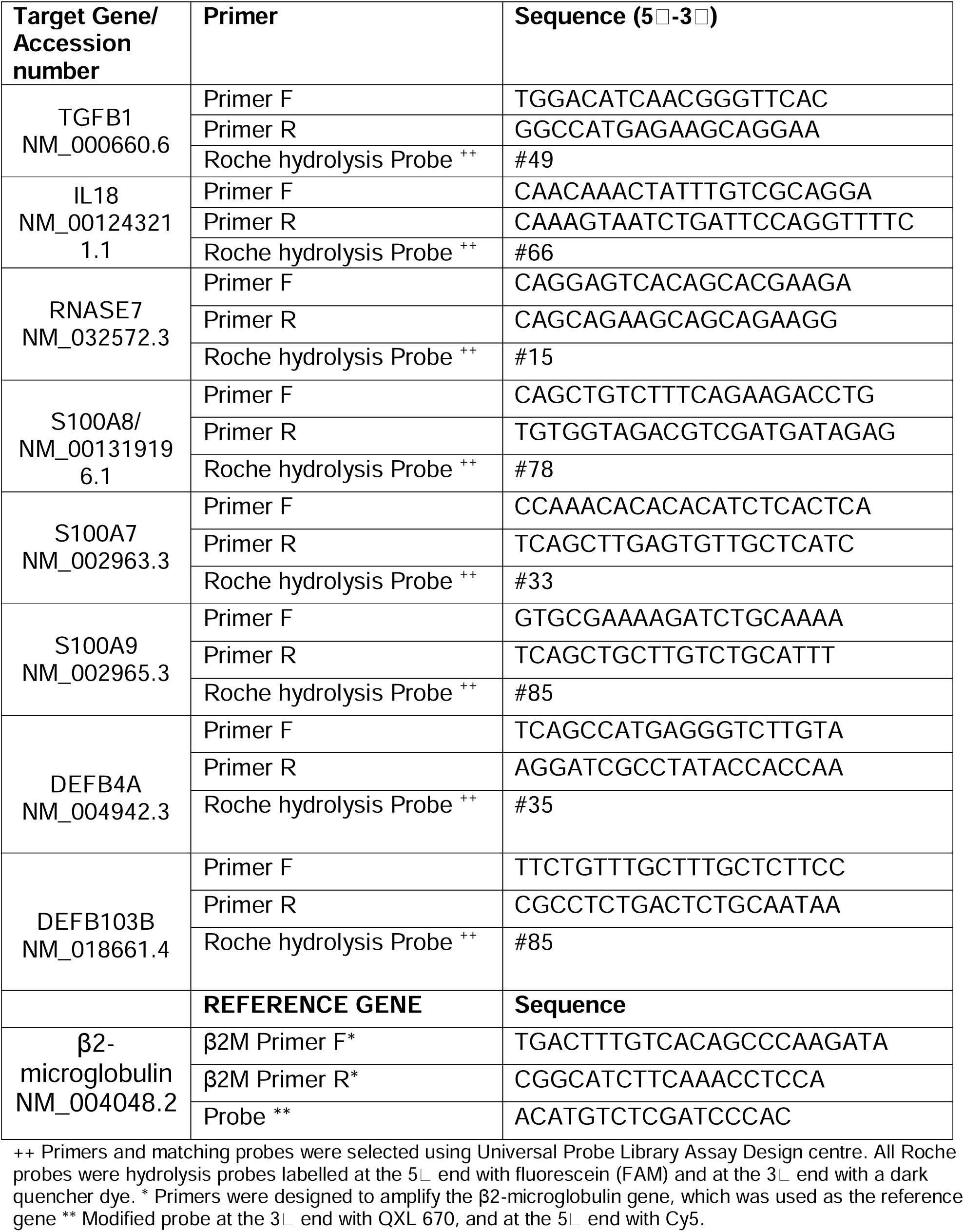
Primers targeting human genes.

**Table 2.** qRT-PCR reactions for skin samples.

| Reagent | Volume (µl) | Final concentration (nM) |
| --- | --- | --- |
| Master mix (Roche) | 5.0 |  |
| 10 µM Target primer forward | 0.25 | 250 |
| 10 µM Target primer reverse | 0.25 | 250 |
| 1 µM Target probe (FAM) | 0.5 | 50 |
| 10 µM Reference primer forward | 0.25 | 250 |
| 10 µM Reference primer reverse | 0.25 | 250 |
| 1 µM Reference probe (Cy5)** | 0.25 | 50 |
| Water | 1.25 | - |
| cDNA | 2.0 | 150-200 ng/µl |
| **Total volume** | **10 µl** |  |

For skin samples, dual hydrolysis probe assays were analysed in the same well, with a FAM probe used for target genes primer pairs and a Cy5 probe for the reference gene.

Reactions were run in a LightCycler 480 (Roche). Following manufacturer’s recommendations, reaction settings were as follows: one cycle at 95 °C for 10 min (ramp rate 4.8 °C/s), 55 cycles of amplification phase with denaturation at 95 °C for 10 s (ramp rate 4.8 °C/s), annealing at 60 °C for 30 s (ramp rate 2.5 °C/s), extension 72 °C for 1 s (ramp rate 4.8 °C/s); and, finally, one cycle of cooling phase at 40 °C for 30 s (ramp rate 2.5 °C/s). Results obtained for each target gene were normalised against β2-microglobulin gene expression levels. The corresponding uninfected/non-inoculated skin samples (with or without OA) were used as negative controls for infection and to measure baseline gene expression levels. Results were analysed using the 2-ΔΔCT method (Livak & Schmittgen, 2001). Statistical analysis was performed using the Student’s t-test or Mann-Whitney test depending on the data distribution using Prism 8 software (GraphPad, La Jolla, CA, USA).

Protein concentration was determined by Coomassie G-250 Bradford protein assay kit following manufacturer’s instructions (Thermo-Fisher Scientific). Protein samples were sent for trypsin digestion and LC-MS/MS analysis (Aberdeen Proteomics Core Facility (www.abdn.ac.uk/ims/facilities/proteomics/)). Four biological replicates were analysed per condition (infected and uninfected skin). Only proteins having two or more identified peptides and two or more Peptide Spectrum Matches (PSM) were selected. Finally, proteins found in at least two out of four analysed samples per condition were included for further Gene Ontology (GO) analysis using the GO consortium online tool (geneontology.org). Area under the curve (AUC) values for each protein were averaged and compared between conditions, inoculated and non-inoculated skin, non-inoculated skin with and without oleic acid and, finally analysed using the Student’s t-test or Mann-Whitney test depending on the distribution of the data, with a value of p<0.05 considered statistically significant (Prism 8 software).

Proteins identified as significant were compared to the CRAPome database (www.crapome.org/). The CRAPome web tool is a Contaminant Repository for Affinity Purification and contains lists of proteins identified in negative control samples, collected using affinity purification followed by mass spectrometry (AP-MS). Proteins found in the CRAPome database in >10% of the cases were not considered significant as they are probably the result of carryover contamination during mass spectrometry experiments.

### Immunoassays

Recovered supernatants from inoculated skin and non-inoculated skin controls were analysed for different cytokines (TGFβ1, TNFα, IL-1β, IL-6, IL-8, IL-18, IFNγ and, TNFRI) at 2-3 days of incubation. Four biological experiments were analysed in duplicate. Multiplex immunoassays were carried out following the manufacturer’s instructions (Milliplex ® Map kits. EMD Millipore Corporation, Livingston UK). Data were analysed using either one-way ANOVA or Kruskal-Wallis test depending on the homogeneity of variance (tested by Bartlett’s test). A *p* value <0.05 was considered statistically significant (Bonferroni correction) and post-hoc analysis done by Dunnett’s or Dunn’s test (Graphpad Prism 8).

## RESULTS

### *M. sympodialis* invaded *ex vivo* skin supplemented with oleic acid and interacted directly with the inner epidermal layer

Host-pathogen interactions between human skin and *M. sympodialis* were investigated using an explant human skin model with skin collected from four healthy donors undergoing cosmetic surgeries (Corzo-Leon et al., 2019). All skin samples were gently wounded by scratching the surface with a needle. Four different skin conditions were used: untreated skin left uninfected or inoculated with *M. sympodialis* (see below) or skin supplemented with 10 µl of 100% oleic acid (OA) and uninfected or inoculated with *M. sympodialis* (MS). Inoculated skin samples were inoculated with 1×10^6^ *M. sympodialis* yeast cells in 10 µl applied onto the skin surface.

Skin inoculated with yeasts without OA supplement did not show macroscopic differences when compared to the non-inoculated skin control after six days. The inoculated skin did not show any macroscopic changes and will be referred to as inoculated skin instead of infected skin. Meanwhile, skin inoculated with yeasts and supplemented with OA had visible yeast growth (Figure 1).

**Figure 1.**
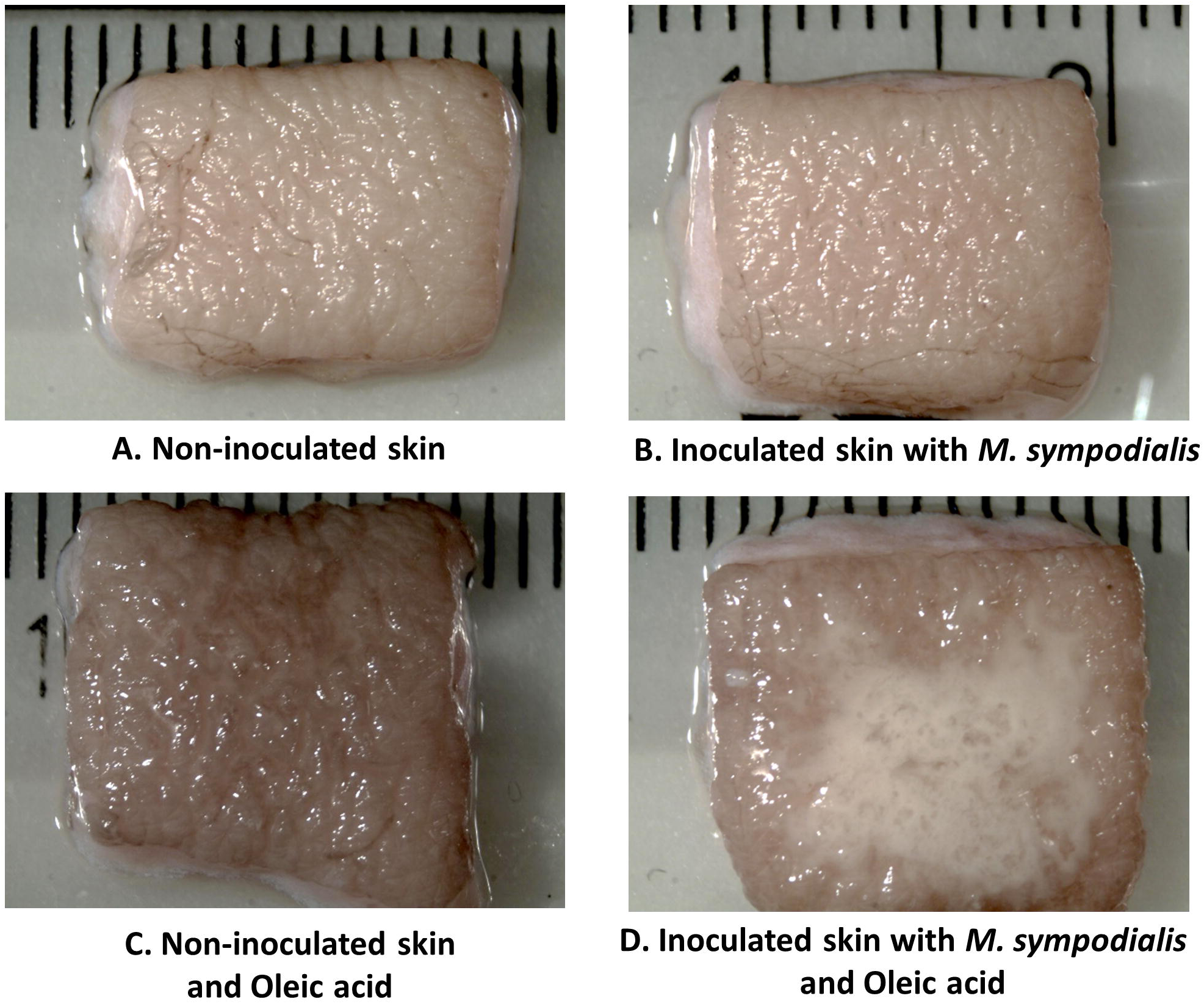
Macroscopic appearance of skin in four different conditions. Human skin explant samples were untreated (A, B) or supplemented with oleic acid (C, D). *M. sympodialis* (1 x 10^6^ yeasts) was inoculated in b & d, with a & c uninoculated. All samples were incubated at 37 □C in 5% CO_2_ for 6 days. Scale is indicated by the ruler, with the space between each bar = 1 mm.

When analysed microscopically by SEM, only the *M. sympodialis*-inoculated skin had fungal structures on the epidermis. The fungal structures differed between the skin supplemented with OA and skin that did not receive OA. Skin without OA (MS or *Malassezia* only) had yeast cells on the top of the skin, while skin receiving both yeasts and OA (MSOA) had not only more abundant yeasts but also elongated fungal structures (Figure 2).

**Figure 2.**
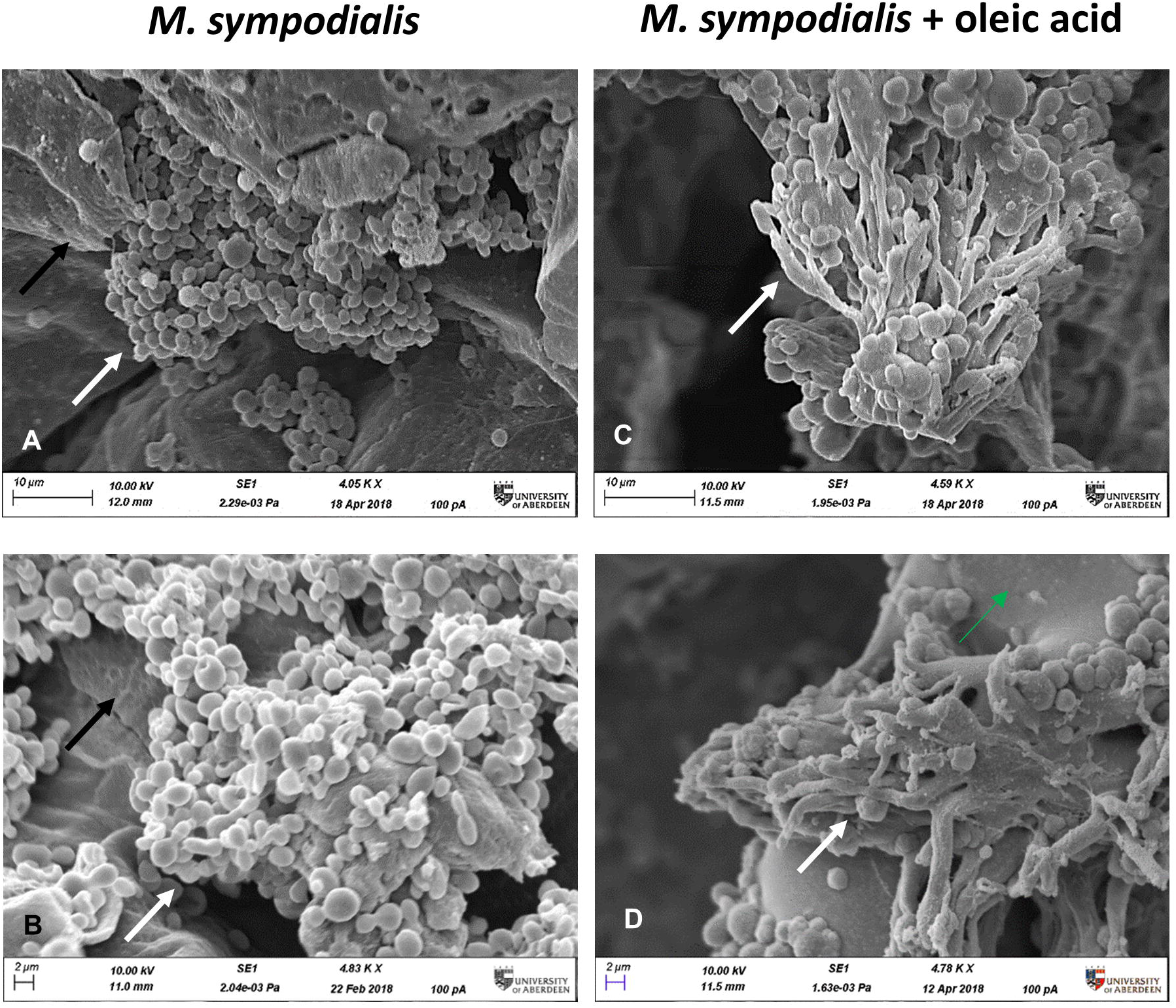
Microscopic appearance of skin inoculated with *M. sympodialis*. *M. sympodialis* yeast cells (1 x 10^6^) were inoculated onto the human skin explants, with or without OA supplementation. All samples were incubated at 37 □C in 5% CO_2_ for 6 days. (A, B) SEM images *M. sympodialis*-inoculated skin, (C, D) skin co-inoculated with *M. sympodialis* and oleic acid. Black arrows indicate corneocytes (skin), white arrows indicate fungal structures. Scale bars represent 10 µm (a, c) or 2 µm (b, d). Images are representative of four biological replicates.

Histological sections, stained with propidium iodide to stain nuclei in the epidermis and calcofluor white (CFW) to stain chitin in the yeast cell walls, were examined to investigate the integrity of keratinocytes and presence of fungal elements, respectively. The epidermis in different samples showed differences in structure. Sections of MSOA skin had detached keratinocyte layers and had yeast cells in direct contact with the inner epidermal layers. In contrast, MS skin had completely intact epidermis and yeast cells were trapped in the outer *stratum corneum* layer. Intact epidermis was also seen in the uninfected skin control without oleic acid. In uninfected skin receiving OA, thinner epidermis was observed but no detachment of epidermal layers was observed (Figure 3).

**Figure 3.**
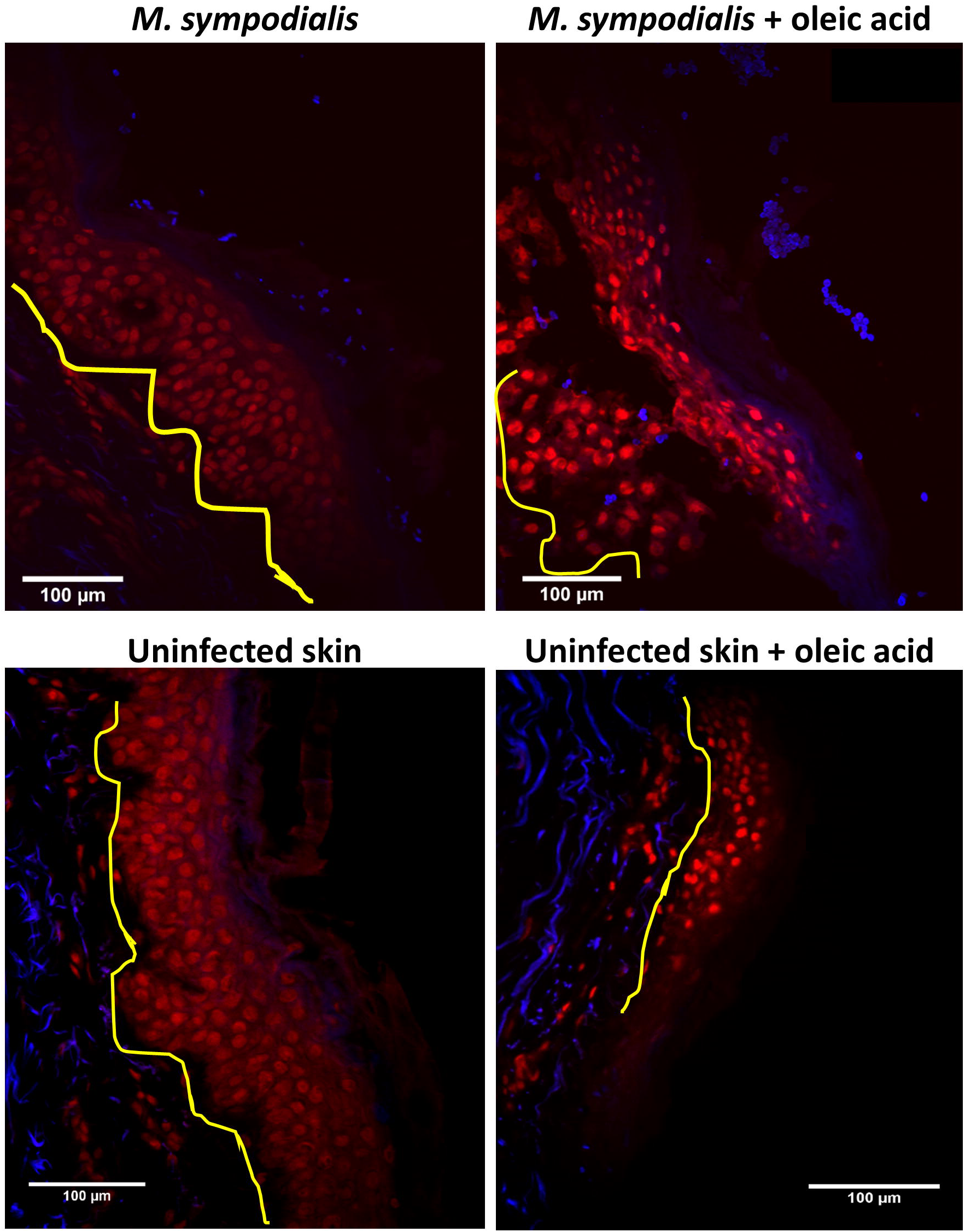
Skin inoculated with *M. sympodialis* with and without oleic acid. Propidium iodide (PI) (red) indicating keratinocytes in epidermis and CFW (blue) staining *M. sympodialis* yeast cells after 6 days incubation. Channels used were DAPI (358 nm/461 nm) for CFW, and Rh-TRITC (543 nm/569 nm) for PI. Yellow line separates dermis from epidermis. Yeasts are seen on the external side of the skin. Images representative of four biological replicates. Scale bars represent 100 µm.

### Local skin response to *M. sympodialis* is characterised by higher expression of genes encoding β-defensin 3, ribonuclease 7 and higher levels of S100 proteins

RNA was extracted from skin tissue samples, at day 6 post-inoculation under the four experimental conditions described above, for gene expression analysis. Due to the importance of AMPs and cytokines in the innate immune response, the expression of eight human genes encoding different AMPs and cytokines was analysed by qRT-PCR. AMP genes included *S100A7* (psoriasin), *S100A8, S100A9, DEFB4A* (β-defensin 2), *DEFB103A* (β-defensin 3), *RNASE7* (ribonuclease 7, RNase7), *IL18*, and *TGFB1* (transforming growth factor β1). Expression of AMP genes was found only in uninfected skin and MS skin (inoculated skin without OA). No expression of AMP genes was found in samples receiving OA. We can confirm that reference genes were amplified and expressed in samples receiving OA verifying that the lack of expression of AMP genes was not due to a technical problem. In MS skin, expression of *DEFB103A* (*p*≤0.03) and *RNASE7* (*p*≤0.03) was increased 193 (IQR 139 to 843) and 7 (IQR 2 to 343) times, respectively, compared to non-inoculated, negative control skin (Figure 4). Meanwhile, *S100A9* was significantly (*p*≤0.03) decreased -10.2 (IQR -15 to -4) fold.

**Figure 4.**
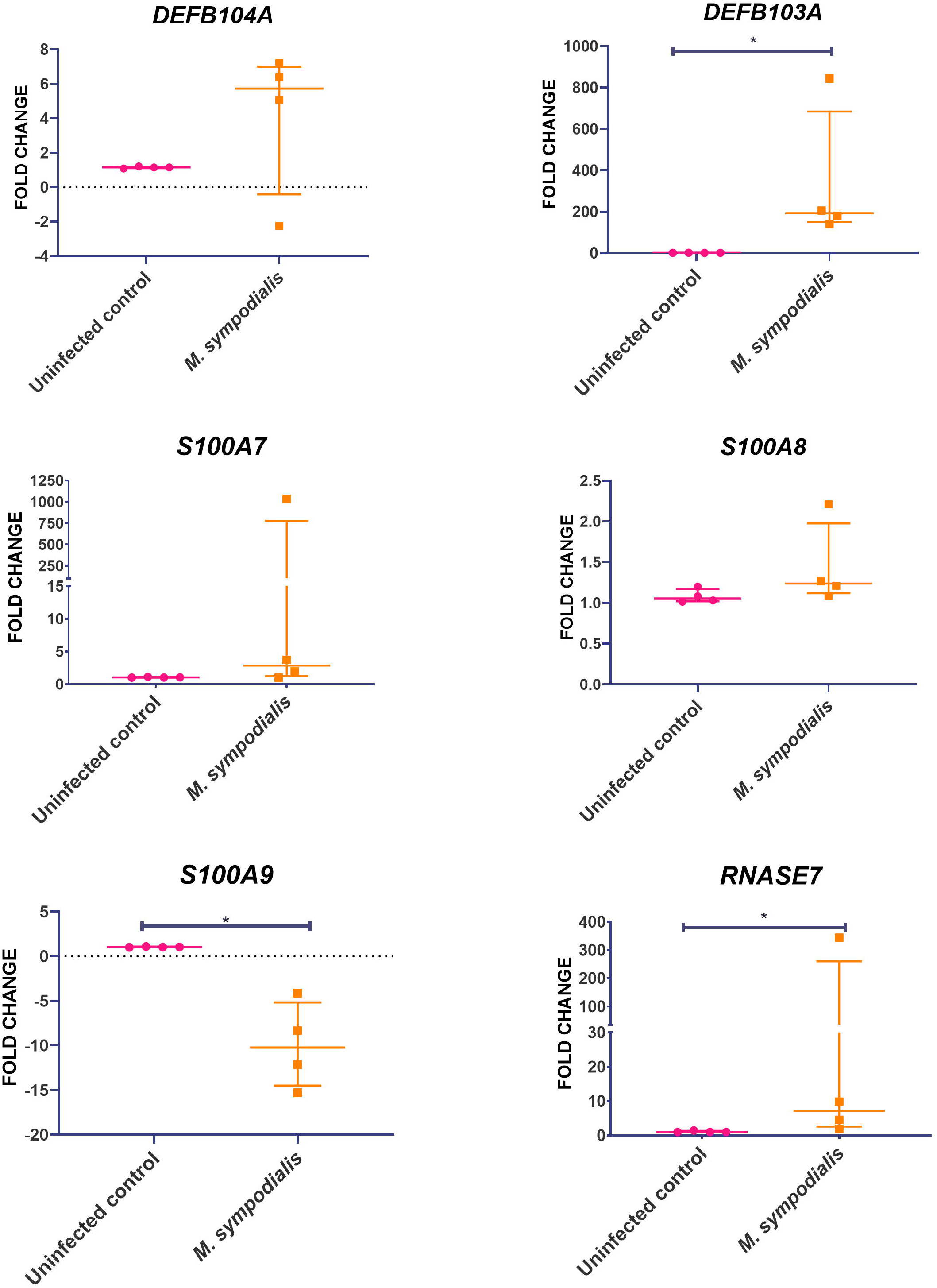
AMP gene expression in *M. sympodialis* infected human skin without OA. RNA was extracted from skin samples after 6 days incubation for qRT-PCR analysis. Results obtained for each target gene were normalised against β2-microglobulin gene expression levels and expressed relative to uninfected skin negative controls. Data is shown as median and IQR, and compared by Mann-Whitney U test. n=4 biological replicates, each analysed in triplicate. \**p* value = ≤0.05

The four different experimental conditions (described above) were also analysed by proteomics at 6 days post-infection to gain a better understanding of the host response to oleic acid and *M. sympodialis* and to investigate the *M. sympodialis* proteome when it was interacting with human skin. Trypsin digestion and protein identification by LC-MS/MS was performed by the Aberdeen Proteomics Facility (Results database in supplementary material S1). Database searches were conducted with the Mascot server v 2.5 using *Homo sapiens* and *M. sympodialis* protein sequences (Swiss-Prot database). Data are available via ProteomeXchange with identifier PXD018404.

A total of 1488 proteins were found in skin tissue of the four biological experiments at the 6 day-time point. After screening for proteins having ≥2 PSM and ≥2 peptides, the total number of proteins included in the analysis was 1048. Non-inoculated skin conditions (with and without OA) were compared. Most of the proteins (98%) found in the non-inoculated skin receiving OA had significantly decreased levels (−2 fold change) when compared to proteins in the non-OA non-inoculated skin control. Fifteen proteins were significantly decreased (*p*≤0.05) in non-inoculated skin receiving OA compared with the non-OA control skin. Five of these proteins are involved in cornification, keratinocyte differentiation and wound healing (KRT5, KRT6A, KRT6B, KRT6C, KRT84) (Supplementary material S3).

Compared to non-inoculated skin, 267/1488 (18%) proteins were greatly increased in the MS condition, while 376/1488 (25%) proteins were greatly increased in the MSOA compared to non-inoculated skin that received oleic acid only. Proteins with higher levels compared to their corresponding non-inoculated control (±OA) were analysed by gene ontology. The most significantly enhanced biological processes in MS skin were cornification (fold change 27, FDR <0.0001), antimicrobial immune response (fold change 16.5, FDR <0.0001), and defence response to fungus (fold change 14, FDR 0.009). Meanwhile, for MSOA, chylomicron remodelling (fold change 38, FDR <0.0001) and removal of superoxide radicals (fold change 32, FDR <0.0001) were the most significantly increased processes (Figure 5). When analysed separately, the proteins in MSOA with significantly increased levels were keratins, including those found decreased in the non-inoculated control with oleic acid (Figure 5, supplementary material S3).

**Figure 5.**
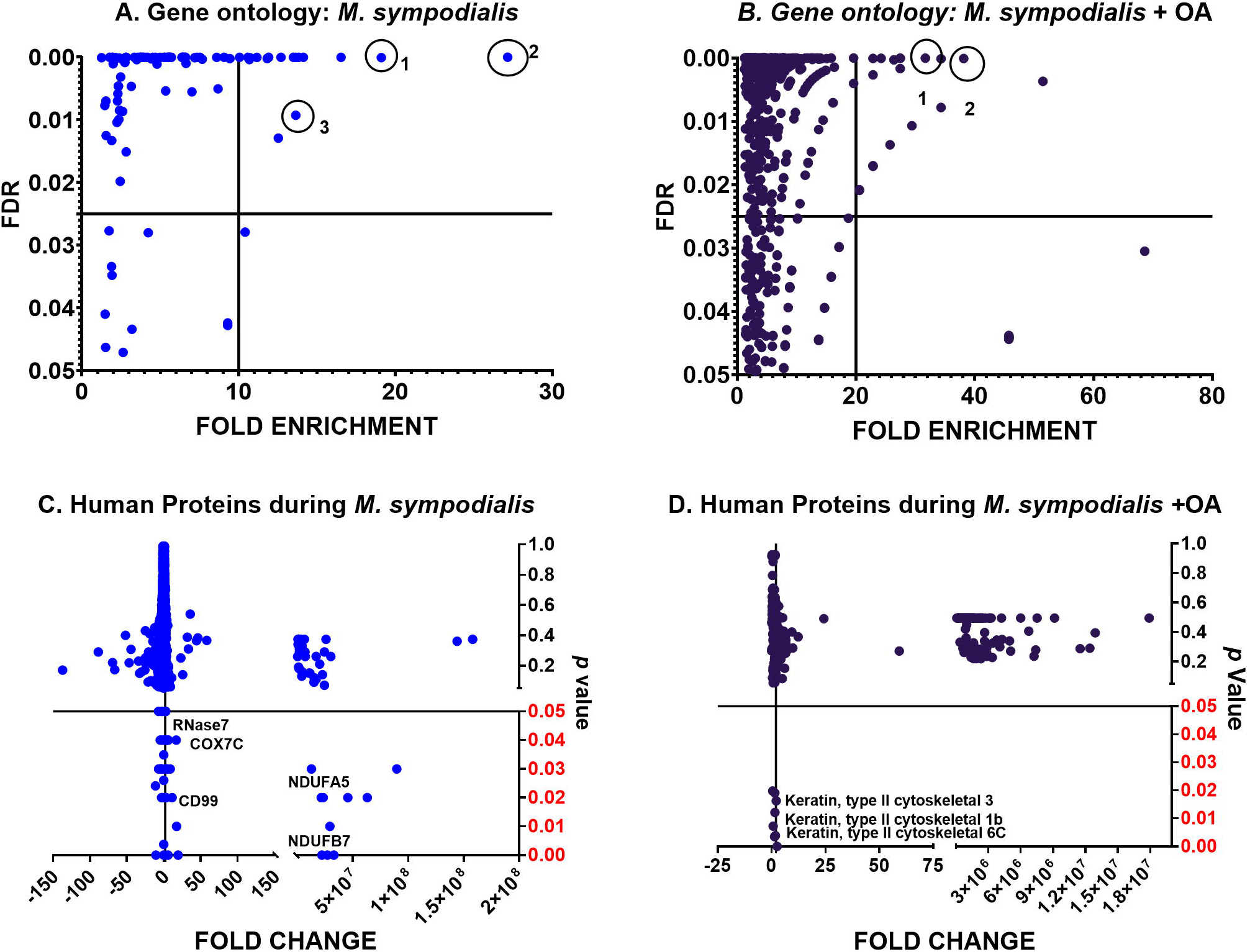
Analysis of human proteins in skin tissue inoculated with *M. sympodialis*. Human proteins with higher levels in tissue at day 6 post-infection (≥2 fold change compared to their corresponding non-inoculated control: non-OA, OA) were analysed by gene ontology. Statistical analysis was done by Fisher’s test with FDR correction (*p* value). (A) biological processes in *M. sympodialis*-inoculated skin, circles are 1. antimicrobial response, 2. cornification, 3. defence response to fungus. (B) processes in skin co-inoculated with *M. sympodialis* and oleic acid, circles are 1. removal of superoxide radicals, 2. chylomicron remodelling. (C) and (D) show proteins with significantly increased levels in each condition. Fold change was estimated relative to protein level in non-inoculated skin without OA.

AMP levels were analysed separately and compared to non-inoculated control by Kruskal-Wallis test and Dunn’s test, as well as using the CRAPome online tool. RNase7 (fold change 2, IQR 1-3, CRAPome 0%, *p*=0.05), S100B (fold change 5, IQR 3-9, CRAPome 0, *p*=0.03), S100A4 (fold change 5, IQR 4-5, CRAPome 2%, *p*=0.01), and S100A2 (fold change 14, IQR 7-24, CRAPome 1%, p=0.05) were significantly increased in the MS skin but not in MSOA skin, where they were either decreased or undetectable (Figure 6). Along with these AMPs, CD99/MIC2, a glycosylated transmembrane protein (fold change 2, IQR 1.7-3.6, *p*=0.05, CRAPome 1%) was also found at higher levels in the MS skin (Figure 6).

**Figure 6.**
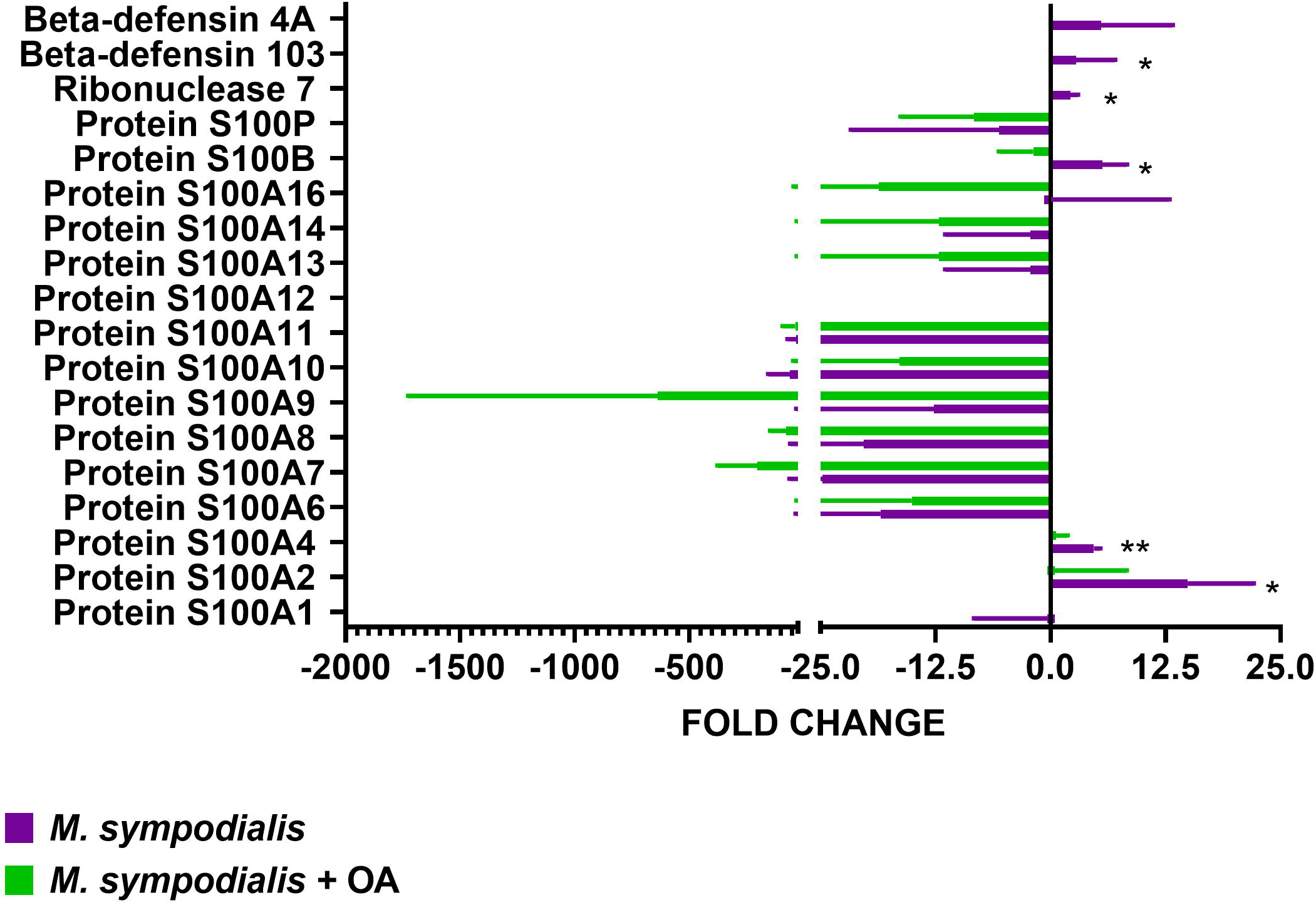
Proteomic analysis of AMP response in skin inoculated with *M. sympodialis*. Single proteins were analysed separately by Kruskal-Wallis test. If *p* value <0.05 after Kruskal-Wallis then post-hoc analysis by Dunn’s test was performed and indicated in the graphic as\**p*≤0.05, ** *p*≤0.001. Data are presented as median and IQR, n=4 biological replicates.

### Secreted cytokine responses of human skin tissue exposed to *M. sympodialis* is characterised by high levels of TGFB1 and IL-18

After two days of incubation supernatants were recovered from the four experimental conditions and analysed for seven different cytokines (IL1-β, IL-6, IL-8, TNFR1, TNFα, TGF1β, IL-18) and compared to non-inoculated skin as the negative control. IL-6 (*p*=0.01) and TNFα (*p*=0.01) showed significant differences across all four conditions, but no difference was found for IL1-β, IL-8, and TNFR1 levels (Figure 7). Post-hoc analysis showed that although IL-6 was significantly (*p*=0.02) different between MS (fold change 1.2, IQR -1.03 to 2.9) and MSOA (fold change -4.4, IQR -9 to -1.1) supernatants, the levels were not different to the negative control. TNFα levels were not different between non-inoculated control and MS skin but were significantly decreased in MSOA skin (fold change -19, IQR -33.4 to -13.7, *p*=0.05) (Figure 7).

**Figure 7.**
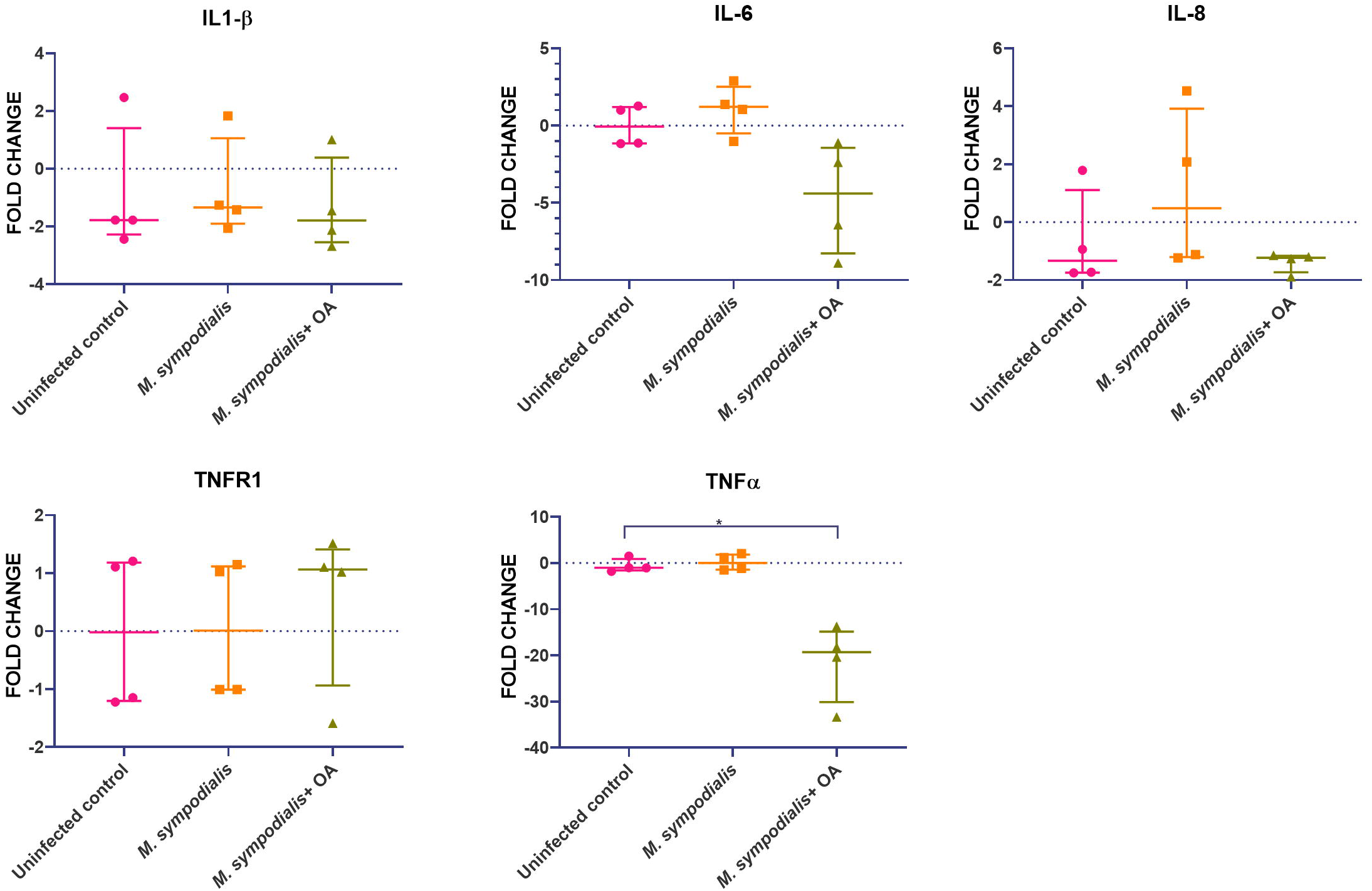
Supernatant cytokine levels from *M. sympodialis*-inoculated skin at two days incubation. Cytokine levels were measured by immunoassay. Fold change was calculated by comparing levels to negative control, non-inoculated skin. Data is shown as median and IQR, and analysed by Kruskal-Wallis test. If *p* value <0.05 after Kruskal-Wallis test then post-hoc analysis by Dunn’s test was performed and indicated in the graphic as \**p*≤0.05. n=4 biological replicates, each analysed in triplicate.

To further explore the role of IL-18 and TGF β1 a more detailed analysis of these two cytokines in tissue and supernatant samples was performed. Gene expression of *TGFB1* (fold change 4, IQR 1.3 to 8.8, *p*=0.001) and *IL18* (fold change 11,805, IQR 8729 to 14495, *p*=0.004) in tissue was increased in MSOA skin, compared to MS skin and non-inoculated skin levels, respectively (Figure 8). Immunoassays were performed to examine levels of TGFβ1 and IL-18 cytokines in supernatants. IL-18 levels increased (fold change 5.6, IQR 4.8 to 10.6, *p*=0.02) in supernatant samples from MSOA skin but not from MS skin (Figure 8). TGFβ1 levels did not change in supernatant.

**Figure 8.**
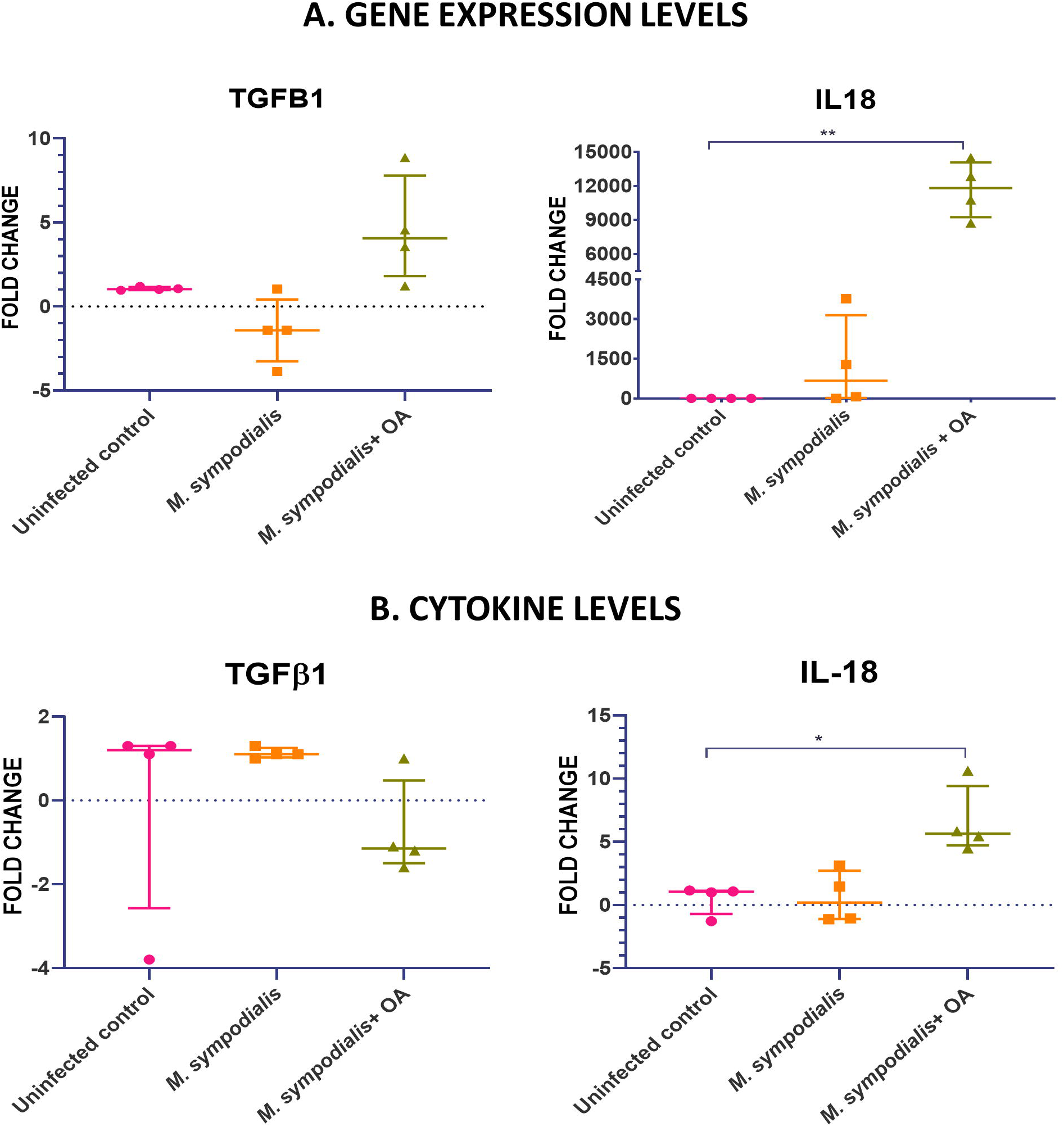
Comparison of TGFB1 and IL-18 expression in tissue and supernatants. For gene expression relative quantification was performed by qRT-PCR using RNA extracted from samples at 6 days post inoculation. Results obtained for each target gene were normalised against β2-microglobulin gene expression levels. Protein levels in supernatant at 2 days post inoculation were measured by immunoassay. In both cases, fold change was estimated relative to expression in non-inoculated skin. Data is shown as median and IQR and statistical analysis was done with Kruskal-Wallis test. If *p* value <0.05 after Kruskal-Wallis test then post-hoc analysis by Dunn’s test was performed and indicated in the graphic as \**p*≤0.05, ** *p*≤0.001. n=4 biological replicates, each analysed in triplicate.

### Nine allergens of *M. sympodialis* were identified in inoculated skin

We next examined protein levels of *M. sympodialis* allergens when the yeast was inoculated onto the skin explant model and incubated for 6 days. A total of 202 *M. sympodialis* proteins were detected by mass spectrometry and identified by matching peptide fingerprints to *M. sympodialis* (reference strain ATCC 42132) protein sequences (Swiss-Prot database). However, after filtering to include proteins found only in *M. sympodialis* inoculated skin in ≥ 2 biological experiments, only 48 fungal proteins were included in the final analysis.

Most of the proteins were found only in MSOA skin (32/48, 66%); 12 (25%) were uncharacterised proteins, 9 (19%) were allergens, and only one lipolytic enzyme, Lipase 3, was identified in MSOA skin. The allergens found in infected skin were Mala S1, Mala S4, Mala S5, Mala S6, Mala S8, Mala S9, Mala S11, Mala S12, Mala S13. Mala S11 and Mala S13 were the only two allergens found in both MSOA and MS skin (Results database in supplementary material S2).

## DISCUSSION

We have characterised the host-pathogen interactions of *M. sympodialis* with human skin at the molecular level using a human skin explant model. The model consisted of inoculating skin explants with *M. sympodialis* yeasts and incubating them for 6 days. The influence of added exogenous oleic acid, mimicking lipid-rich skin niches, was also investigated. First, the presence of yeasts were confirmed by fluorescence microscopy and SEM in the inoculated samples. Then, host responses were evaluated by gene expression, proteomics and immunoassays.

In this model, increased mRNA and protein expression of β-defensin 3 and RNase 7 was detected in MS skin, similar to previous reports in skin of individuals with pityriasis versicolour and atopic dermatitis (Brasch *et al*, 2014; Gambichler *et al*, 2008). Meanwhile, AMP gene expression was not detected in OA or MSOA skin. Oleic acid conditions were added to this model to evaluate the host response to *M. sympodialis* in oily skin conditions. Previous studies found fatty acids (oleic acid, palmitic acid and lauric acid) to be protective, resulting in the upregulation of β-defensin 2 expression by sebocytes in cell culture (Nakatsuji *et al*, 2010). This is contrary to what was observed with this model in non-inoculated and MSOA skin and may be due to the high concentration of OA used for the experiments in the current report. Skin damage by OA has been documented previously with concentrations as low as 5% and is macroscopically evident as dermatitis in healthy volunteers and histological damage in reconstructed epidermis (Boelsma *et al*, 1996). Therefore the lack of AMP expression in OA or MSOA skin may be due to epidermal damage as a result of OA supplementation and would require further validation. The direct effect of OA on skin was analysed here by proteomics and the majority of proteins (including keratins involved in cornification, keratinocyte differentiation and wound healing responses) were detected at lower levels in OA skin compared to untreated skin. Only a handful of proteins had increased levels in OA skin but the differences in fold change were not statistically significant. It is known that OA can 1) damage and produce desquamation of the *stratum corneum* (DeAngelis *et al*, 2005), which was also observed in histological sections in our model and 2) histological damage facilitates penetration of *M. sympodialis* so that it contacts and damages keratinocytes in deeper skin layers. These two consequences could result in the absence of AMPs produced by the epidermis in MSOA skin.

The lack of AMP protein expression in MSOA skin in this model is similar to what has been reported in skin lesions of AE individuals, where previous studies have documented decreased or no expression of β-defensin 2 and LL37 in acute and chronic skin lesions of AE individuals (Ong *et al*, 2002; Clausen *et al*, 2018). Another report also showed both AMP gene and protein expression in the epidermis was lower in atopic dermatitis compared to psoriasis patients(de Jongh et al, 2005).

High levels of certain S100 proteins were found in MS skin (S100A2, S100A4 and S100B). These have not been reported to have a role in *Malassezia* infections or allergic reactions. However, these three S100 proteins have been reported to have a role in macrophage migration, cell proliferation and migration, and as apoptosis inhibitors and regulators of p53 protein (Donato *et al*, 2013). The increase of these S100 proteins should be confirmed with different techniques, such as gene expression, immunoassays or immunofluorescence as LC-MS/MS could misidentify these peptides with other similar proteins sharing S100 domains such as filaggrin (Bunick *et al*, 2015). This differentiation will be important as filaggrin is essential for epidermal barrier formation (Hänel *et al*, 2013) and the loss of its function is already recognised as a causative factor for AE (Nutten, 2015). Further study of the role of filaggrin in the response to *M. sympodialis* is required. In addition, the lower expression of *S100A9* gene found in skin inoculated with *M. sympodialis* has not been reported before. The expression of *S100A9* gene was expected to be increased or unchanged as seen with the rest of S100 proteins. In order to investigate whether decreased expression of *S100A9* is a feature of *M. sympodialis* skin response, a follow up of the dynamics of *S100A9* gene and protein expression in earlier and later stages of *M. sympodialis* infection would be required.

The effect of *Malassezia* species on cytokine production can vary depending on the clinical and experimental context (Watanabe *et al*, 2001; Pedrosa *et al*, 2019; Faergemann *et al*, 2001). High levels of IL-1β, IL-6, IL-8 and TNFα have been found in supernatants of human keratinocyte cell cultures after 3 to 6 h of co-incubation (Watanabe *et al*, 2001). However, the cytokine response to *Malassezia* sp. varies depending on the species; *M. pachydermitis* induced the highest levels of all these cytokines and *M. furfur* induced almost no response (Watanabe *et al*, 2001). In our study, none of these cytokines had high levels when MS skin was compared to non-inoculated controls and TNFα levels were significantly decreased in MSOA skin. This finding can be explained in three ways. Firstly, the time point for cytokine analysis differs. In a recent study using reconstructed epidermis, gene expression of *IL1B, IGFB1* and *TNFA* in tissue was increased after 6 h incubation with *M. sympodialis*, but these same genes were downregulated after 48 h of co-incubation (Pedrosa *et al*, 2019). Secondly, models to study host response to fungal infection can differ from the response in real human infections. Faergemann *et al*. (2001) reported no difference in skin TNFα levels in individuals with *Malassezia* folliculitis and seborrheic dermatitis when compared to healthy volunteers, similar to the current *ex-vivo* skin model and contrary to what has been reported from monolayer keratinocyte culture. In addition, lower TNFα serum levels were found in individuals with chronic AE, along with low serum levels of IL-10, β-defensin 3 and high levels of β-defensin 2 (Kanda & Watanabe, 2012). Finally, the lack of AMP expression seen in the MSOA skin can explain the low levels of some cytokines as these AMPs (especially, β-defensins, S100 proteins and cathelicidin) are key for inducing cytokine responses (Niyonsaba *et al*, 2017).

*IL18* and *TGFB1* were highly expressed in MSOA skin, whilst only IL-18 levels were significantly higher in the analysed supernatants. IL-18 belongs to the IL-1 cytokine family, along with IL-1α and IL-1β, and is cleaved by caspase-1 after being activated by 3NLRP inflammasome (Fenini *et al*, 2017). This inflammasome pathway is crucial for inducing Th1/Th2 responses. IL-18 induces the formation of high serum levels of IgM, IgG_1_, IgG_2a_ and very high levels of IgE antibodies (Enoksson *et al*, 2011). The production of these antibodies depends on CD4^+^ T cell-derived IL-4 and the auto-reactivity of these antibodies is regulated and depends on NK T cells (Enoksson *et al*, 2011). The production of high levels of IL-18, along with high levels of IL-4 (Th2 biased response), has been associated with worse prognosis in other infections such as leishmaniasis (Gurung *et al*, 2015). High serum levels of IL-18, IL12/p40 and IgE antibodies have been found in individuals with atopic eczema and their serum levels correlate proportionally with clinical severity of AE skin lesions (Zedan *et al*, 2015).

As mentioned previously, *M. sympodialis* allergens play a crucial role in the pathogenesis of atopic dermatitis (Gioti *et al*, 2013). In this study, higher numbers of allergens were identified in the MSOA skin compared to MS skin. This finding could be explained by the higher number of yeasts present on the surface of MSOA skin. Due to the skin damage caused by OA, it is possible that these allergens were in contact with inner epidermal cells and contributed to the host response seen in MSOA skin. The role of allergens in *M. sympodialis* pathogenicity is a future avenue of investigation with this *ex-vivo* human skin model.

In conclusion, the local host response to *M. sympodialis* can be characterised using this *ex-vivo* human skin model. Such host response can vary as previously described, depending on the *Malassezia* species, host intrinsic and extrinsic factors, time of clinical evolution, and type of infection. Most of these conditions can potentially be mimicked in this *ex-vivo* skin model. Comparison of responses between different skin conditions has already been shown to be possible in this study. In non-oily and intact skin, AMPs and S100 proteins are key in the response to *M. sympodialis*, but in oily and damaged skin, allergens and yeasts are in direct contact with keratinocytes, and inflammasome responses seem to lead to increased IL-18, which can promote chronic inflammation, auto-reactivity in skin and continuing local damage.

## AUTHORS CONTRIBUTION

DECL, DMM and CM conceptualised and designed the study. DECL performed the experiments and data analysis. DECL, DMM and CM wrote the manuscript, reviewed and approved the submitted version.

## ACKNOWLEDGMENTS

Thanks to the Technology hubs at the University of Aberdeen (Microscopy and Histology, qPCR, Proteomics) for their support, sample processing and training. Special thanks to Professor Annika Scheynius from the Karolinska Institute, Stockholm, Sweden for sharing her expertise and constructive discussions and for giving us the inspiration to work on *Malassezia*.

## Supplementary material

S1. Results supplementary material S1 proteomics human database OA

S2. Results supplementary material S2 *Malassazia sympodialis* proteins

S3. Results supplementary material S3 Proteomic analysis of uninfected skin samples with and without OA

